# Effect of field strength on RF power deposition near conductive leads: A simulation study of SAR in DBS lead models during MRI at 1.5 T - 10.5 T

**DOI:** 10.1101/2022.12.13.520303

**Authors:** Ehsan Kazemivalipour, Alireza Sadeghi-Tarakameh, Boris Keil, Yigitcan Eryaman, Ergin Atalar, Laleh Golestanirad

**Affiliations:** Department of Radiology, Feinberg School of Medicine, Northwestern University, Chicago, Illinois, USA; Electrical and Electronics Engineering Department, Bilkent University, Ankara, Turkey; National Magnetic Resonance Research Center (UMRAM), Bilkent University, Ankara, Turkey; Athinoula A. Martinos Center for Biomedical Imaging, Department of Radiology, Massachusetts General Hospital, Harvard Medical School, Charlestown, Massachusetts, USA; Harvard Medical School, Boston, Massachusetts, USA; Center for Magnetic Resonance Research (CMRR), University of Minnesota, Minneapolis, Minnesota; Institute of Medical Physics and Radiation Protection, TH Mittelhessen University of Applied Sciences, Giessen, Germany; Department of Biomedical Engineering, McCormick School of Engineering, Northwestern University, Evanston, Illinois, USA

**Keywords:** Deep brain stimulation (DBS), magnetic resonance imaging (MRI), MRI safety, medical implants, specific absorption rate (SAR), ultrahigh field MRI

## Abstract

**Background:** Since the advent of magnetic resonance imaging (MRI) nearly four decades ago, there has been a quest for ever-higher magnetic field strengths. Strong incentives exist to do so, as increasing the magnetic field strength increases the signal-to-noise ratio of images. However, ensuring patient safety becomes more challenging at high and ultrahigh field MRI (i.e., ≥3 T) compared to lower fields. The problem is exacerbated for patients with conductive implants, such as those with deep brain stimulation (DBS) devices, as excessive local heating can occur around implanted lead tips. Despite extensive effort to assess radio frequency (RF) heating of implants during MRI at 1.5 T, a comparative study that systematically examines the effects of field strength and various exposure limits on RF heating is missing.

**Purpose:** This study aims to perform numerical simulations that systematically compare RF power deposition near DBS lead models during MRI at common clinical and ultra-high field strengths, namely 1.5, 3, 7, and 10.5 T. Furthermore, we assess the effects of different exposure constraints on RF power deposition by imposing limits on either the B_1_^+^ or global head specific absorption rate (SAR) as these two exposure limits commonly appear in MRI guidelines.

**Methods:** We created 33 unique DBS lead models based on postoperative computed tomography (CT) images of patients with implanted DBS devices and performed electromagnetic simulations to evaluate the SAR of RF energy in the tissue surrounding lead tips during RF exposure at frequencies ranging from 64 MHz (1.5 T) to 447 MHz (10.5 T). The RF exposure was implemented via realistic MRI RF coil models created based on physical prototypes built in our institutions. We systematically examined the distribution of local SAR at different frequencies with the input coil power adjusted to either limit the B_1_^+^ or the global head SAR.

**Results:** The MRI RF coils at higher resonant frequencies generated lower SARs around the lead tips when the global head SAR was constrained. The trend was reversed when the constraint was imposed on B_1_^+^.

**Conclusion:** At higher static fields, MRI is not necessarily more dangerous than at lower fields for patients with conductive leads. Specifically, when a conservative safety criterion, such as constraints on the global SAR, is imposed, coils at a higher resonant frequency tend to generate a lower local SAR around implanted leads due to the decreased B_1_^+^ and, by proxy, **E** field levels.

## 1 INTRODUCTION

Magnetic resonance imaging (MRI) has become a powerful imaging modality in the arsenal of noninvasive diagnostic tools, providing an unparalleled spatial resolution and soft-tissue contrast, as well as allowing to monitor functional changes in tissue. Since the advent of MRI nearly four decades ago, there has been a race toward ever-increasing magnetic fields. There are strong incentives to do so: increasing the strength of the static magnetic field substantially increases the signal-to-noise ratio to visualize small structures previously unobservable on MRI scans. For example, MRI at 7 T can differentiate subsegments of the globus pallidus, a small brain nucleus that is the target of neuromodulation therapies, such as deep brain stimulation (DBS)[1].

With MRI becoming increasingly prevalent, the number of cases in which patients with conductive implants are referred for an MRI exam increases. It is estimated that 50% to 75% of patients with cardiovascular implants or neuromodulation devices will require MRI during their lifetime [2, 3], with many patients requiring repeated examinations [4]. Nevertheless, performing MRI in the presence of electronic implants that typically have elongated conductive leads is still challenging due to the risks associated with the radio frequency (RF) heating of implants. This phenomenon, commonly known as the *antenna effect*, occurs when the electric field of the MRI transmit coil induces current on conductive lead wires, which raises the specific absorption rate (SAR) of the RF energy in the tissue surrounding the lead tips [5-7]. Excessive tissue heating and serious thermal injuries could arise from this mechanism [8]. Therefore, the criteria under which patients with conductive implants are indicated for MRI are restrictive. Most patients with DBS devices, for example, can only undergo MRI at 1.5 T scanners with pulse sequences with substantially reduced power which are not optimal for target visualization [9-11].

Extensive effort has been dedicated to assessing RF heating of elongated implants during MRI at 1.5 T and, to a lesser degree, at 3 T and above [12]. However, a comparative study examining the differential effects of resonant frequency on implant heating in the wide range of currently available MRI systems is missing. This study is particularly important as misconceptions exist among some MRI users that scanners at higher field strengths are inherently more dangerous in terms of local implant heating.

We report the results of a systematic simulation study with 33 patient-derived models of DBS leads exposed to RF radiation in a range of frequencies corresponding to 1.5 to 10.5 T MRI and compare the local SAR around the lead tips for different exposure limits. Because the trajectory of an implanted lead substantially affects local power deposition near the lead’s tip [13-17] and because we aimed to generate data representative of the clinical population, lead models have trajectories reconstructed from computed tomography (CT) images of patients with DBS devices. Moreover, models of MRI RF coils were recreated based on physical prototypes that we have built for neuroimaging studies; thus, their field distributions represent realistic scenarios. We evaluated the maximum of a 1 g-averaged SAR in cubic volumes surrounding the tips of DBS lead models for exposure conditions that put a cap on a) the magnitude of B_1_^+^ and b) the magnitude of the global head SAR (GHSAR). These two exposure constraints were selected as they are commonly found in the MRI-conditional labeling of devices. Older guidelines imposed the limit on the global SAR, which tends to be more conservative, whereas recent guidelines impose limits on B_1_^+^.

## 2 METHODS

All models described below are available to be used for research upon sending written request to the corresponding author.

### 2.1 RF transmit coils

Coil geometries were based on physical prototypes built in our labs. Four RF transmit head coil models were constructed and tuned to respective proton Larmor frequencies at 1.5, 3, 7, and 10.5 T (64 MHz for 1.5 T, 127 MHz for 3 T, 297 MHz for 7 T, and 447 MHz for 10.5 T). These models included two 16-rung low-pass birdcage coils at 1.5 T [18] and 3 T [19], a 16-rung shielded hybrid birdcage coil at 7 T [20] and an 8-channel bumped dipole array at 10.5 T [21]. Figure 1 presents the geometrical details of the coils loaded with a homogeneous human head model.

**Figure 1:**
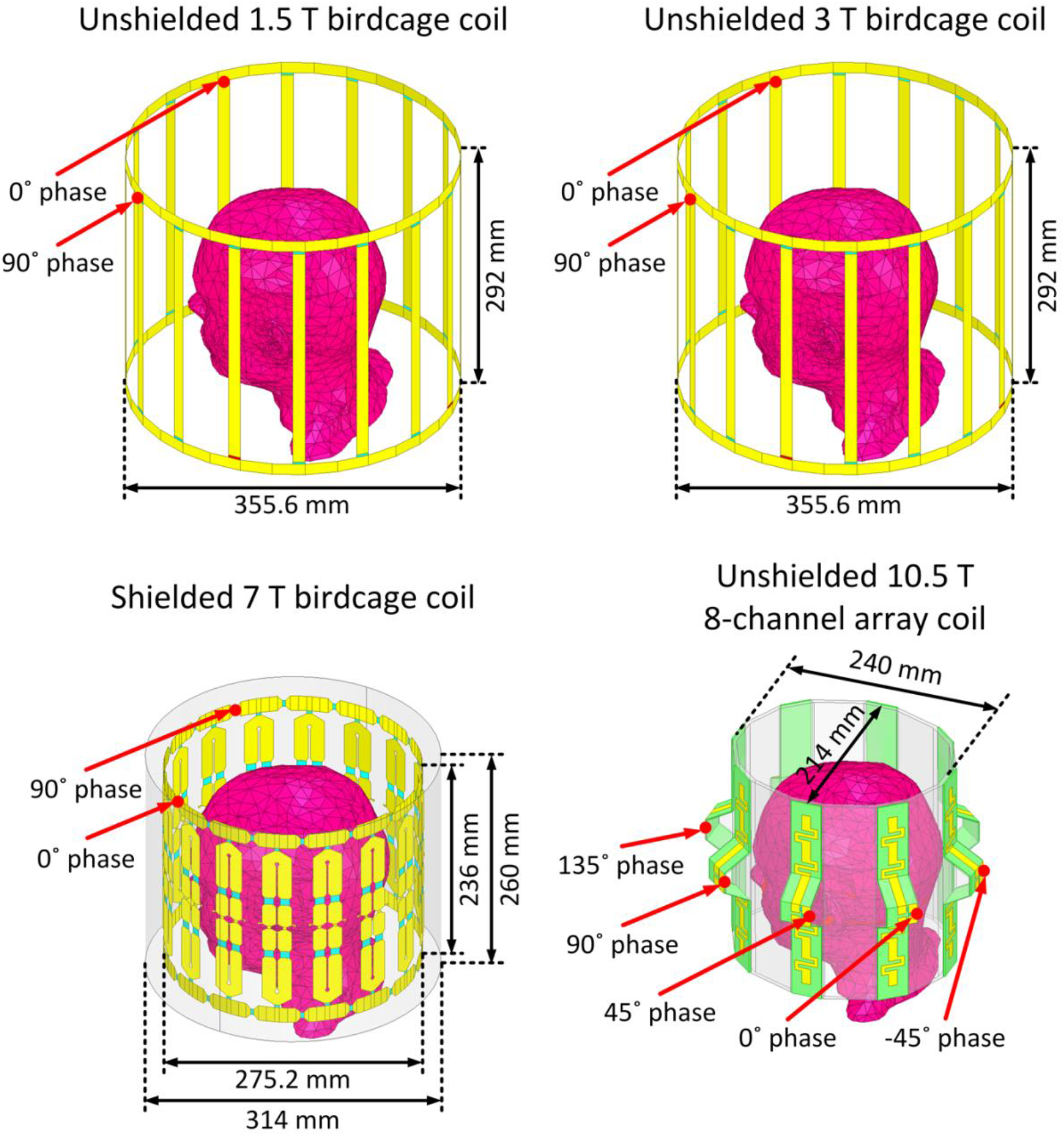
Numerical models of a 1.5 T 16-rung low-pass birdcage coil tuned at 64 MHz, a 3 T 16-rung low-pass birdcage coil tuned at 127 MHz, a 7 T 16-rung hybrid birdcage coil tuned at 297 MHz, and a 10.5 T 8-channel bumped dipole array tuned at 447 MHz all loaded with a homogeneous human head model with no implant. The geometrical dimensions of the coils were the same as the coils reported in literature[18-21].

The 1.5, 3, and 7 T birdcage coils were excited using two ports, each fed with the same amplitude and a 90° phase difference to achieve circular polarization (CP mode) in the sample. The 8-channel bumped dipole 10.5 T coil was excited using eight ports, each fed with the same amplitude and a 45° phase difference to create a CP-like mode. Figure S1 in the supporting information displays B_1_^+^ and **E** field distributions on an axial plane passing through the iso-center of each coil. As observed, the B_1_^+^ uniformity in the CP excitation mode was decreased by increasing the field strength. This outcome was expected due to the lower ratio of the RF wavelength to the body model dimensions at higher field strengths [22, 23].

### 2.2 DBS lead models with realistic trajectories

It is established that the trajectory of an elongated lead and its position within the MRI coil strongly affects its RF heating [24-30]. The extracranial DBS lead trajectories (i.e., the trajectory of the portion of the lead that is outside of the brain and placed over the skull) vary substantially from patient to patient, depending on the surgeon’s practice styles. Thus, studies that aim to assess RF heating of DBS implants should ideally include clinically relevant lead trajectories reflective of the heterogeneity observed in the patient population. In this study, we simulated 33 unique DBS lead models with realistic patient-derived trajectories based on postoperative CT images of 20 patients operated in our institutions. Use of patient data for the purpose of MRI RF heating assessment and publication of de-identified images was approved by Northwestern University’s Institutional Review Board (STU00206859). Consent was waived as the data was collected retrospectively and analyzed anonymously. The details of image segmentation and model creation were similar to our previous studies [13, 31-33].

Out of the 33 simulated lead models, 26 models were constructed based on the CT images of patients with bilateral leads (ID1 – ID13, Figure 2) and 7 models were constructed based on the CT images of patients with unilateral leads (ID14 – ID20, Figure 2). Each lead model comprised four electrode contacts connected via a solid straight platinum core (diameter = 0.26 mm and *σ* = 9.3×10^6^ S/m), embedded in a urethane insulator (diameter = 1.27 mm and *ε*_*r*_ = 3.5). All lead models were 40 cm long. Lead models were incorporated into a homogeneous human head model with the properties of average tissue (σ = 0.49 S/m and *ε*_*r*_ = 66). Note that our models represent lead-only DBS systems, that is, extension cables (∼ 60 cm) and the pulse generator were not included. Also, note that we chose to keep the dielectric properties of the issue constant over the range of studied frequencies to reduce the number of factors that were varied among the coils.

**Figure 2:**
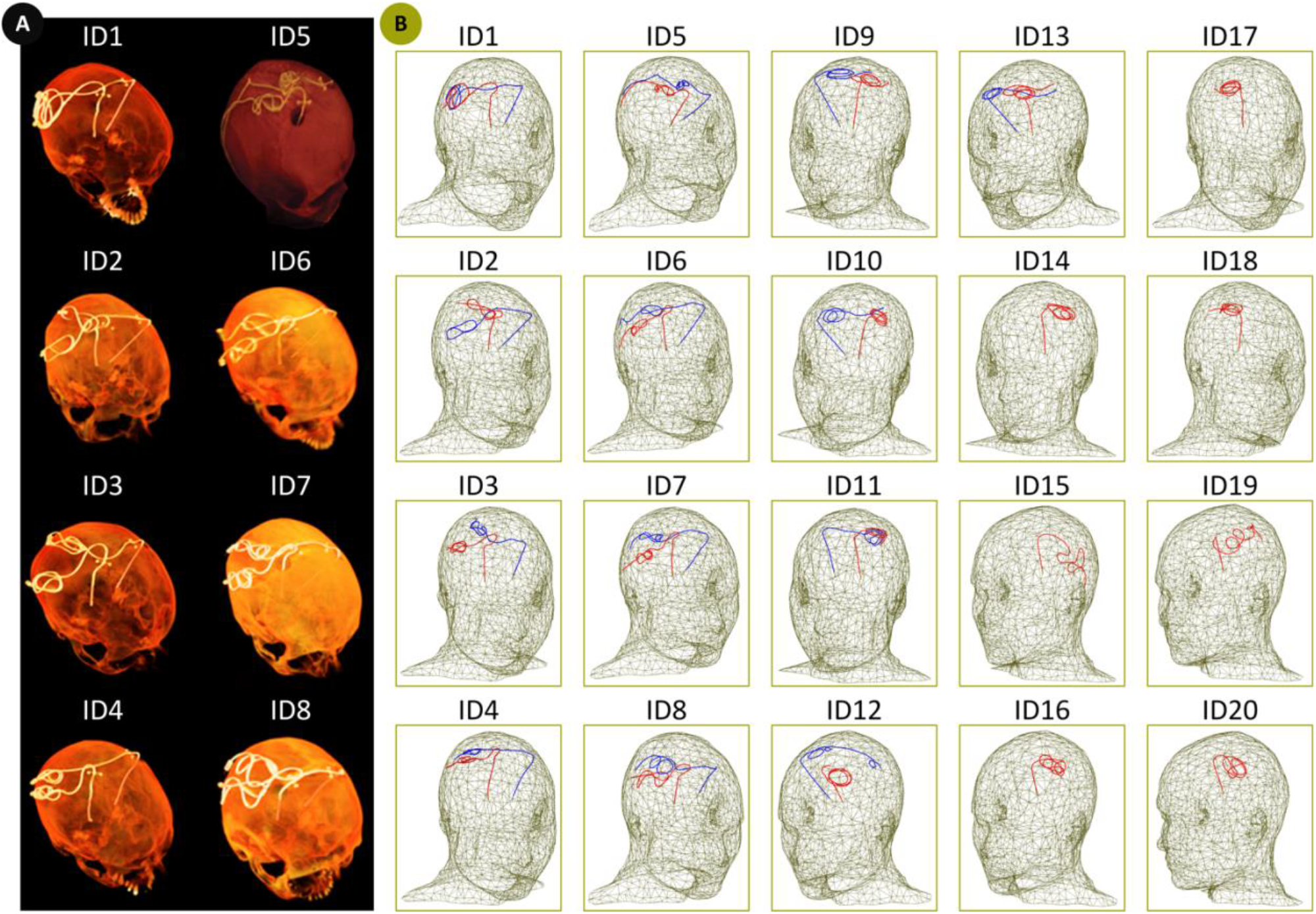
(A) Examples of postoperative CT images (ID1-ID8). (B) Reconstructed models of isolated DBS leads (ID1 - ID20). Lead trajectories were constructed based on the postoperative CT images of 13 patients with bilateral DBS implantations (ID1 - ID13), and 7 patients with unilateral DBS implantations (ID14 - ID20) and were incorporated in a homogeneous head model for EM simulations.

### 2.3 Numerical simulation

Electromagnetic (EM) simulations were implemented in ANSYS Electronics Desktop 2019 R2 (ANSYS Inc., Canonsburg, PA) on a Dell PowerEdge R740xd server equipped with two Intel(R) Xeon(R) Gold 6140 2.3 GHz processors and 1.5 TB of RAM. The EM fields induced in the head and lead models by each coil were computed using a numerical method based on the combined FEM simulation and circuit analysis [13, 34-38]. The maximum of 1 g-averaged SAR (1 g-SAR_max_) was calculated within a high mesh resolution cubic volume of 20 × 20 × 20 mm surrounding the DBS lead contacts (referred to as the SAR box). As a reminder, the point-wise local SAR is defined as follows:

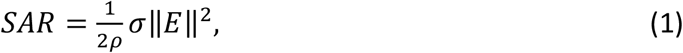

where ‖*E*‖ denotes the magnitude of the electric field (V/m), *ρ* is the mass density of the material (kg/m^3^), and σ represents the material conductivity (S/m). The averaged SAR is calculated over a volume that surrounds each mesh point. The volume used in the 1 g estimation is determined at each mesh point by a closed surface with a volume such that the sum of the product of each material subvolume within totals 1 g by its mass density.

For each coil, we adjusted the input power to impose limits on 1) the GHSAR (3 W/kg) and (2) spatial average of B_1_^+^ on an axial plane passing through the center of the head (B_1_^+^= 2 μT). These two outcome measures were chosen because they are automatically calculated and reported by the scanner and are commonly adopted in MRI-conditional guidelines. The plane over which B_1_^+^was sampled was 40 mm below the coil isocenters, allowing unbiased B_1_^+^ averaging without implant-induced perturbations (Figure 3).For all simulations, the edge length of the tetrahedral mesh elements inside the head model was confined to <10 mm, except for the SAR boxes, which enforced the edge lengths to <1 mm. In addition, the triangular mesh edge lengths on the surfaces of all electrodes and their insulation were restricted to <0.5 mm. Finally, the maximum size of the mesh elements on the RF coil components was set to <5 mm, except for the lumped ports, in which the maximum element length was restricted to <1 mm. The ANSYS high-frequency structure simulator was set to follow an adaptive meshing process, where the mesh was refined until the magnitude of the difference between the scattering parameters calculated in two consecutive iterations was less than 0.01. We demonstrated that the convergence criteria based on the S-parameters also guarantees the convergence of the electric field (and SAR) around the implanted leads.[39] All simulations converged within three adaptive passes. The simulation time was ∼12 hours for each model.

**Figure 3:**
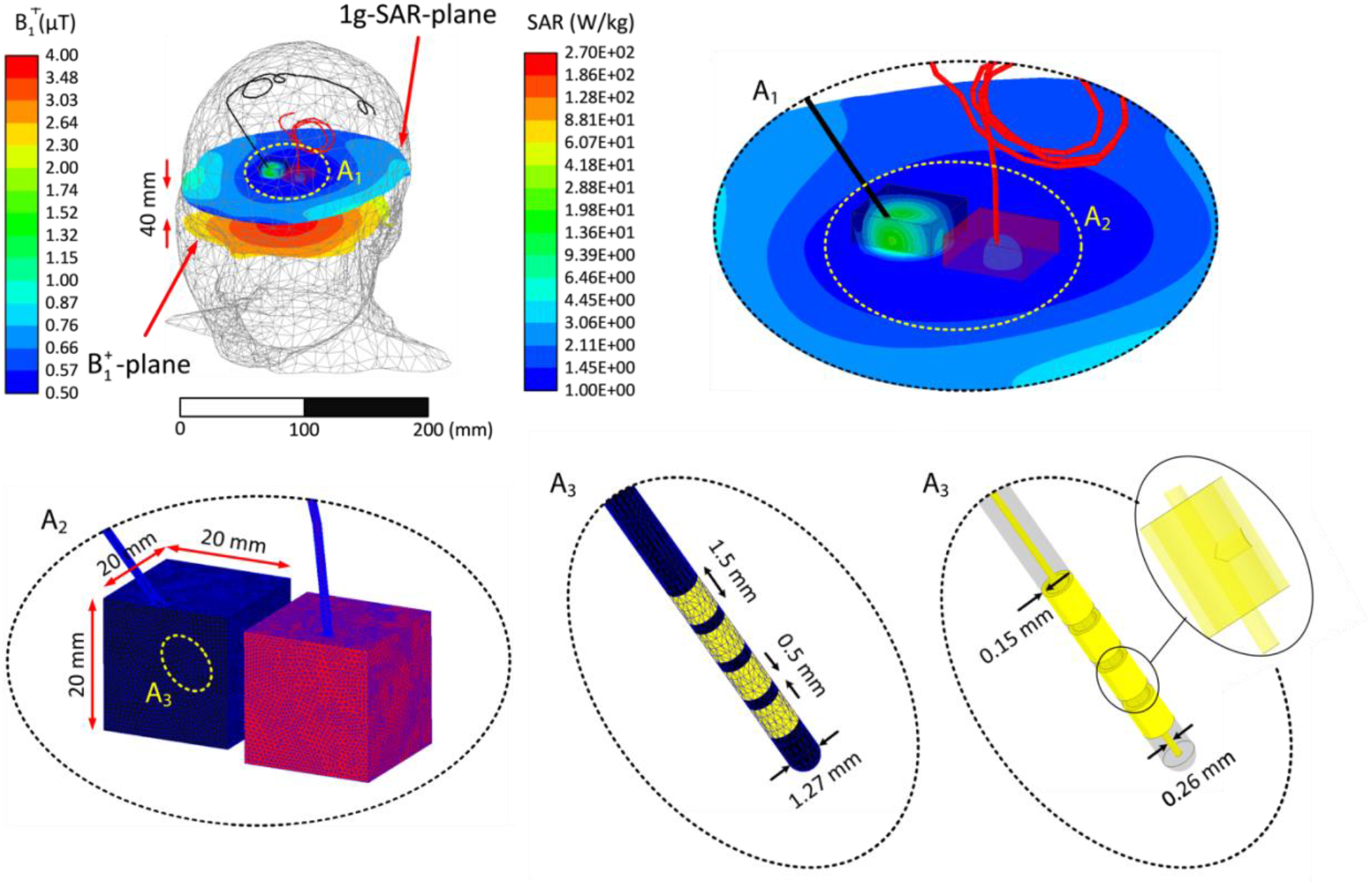
Example of mesh density in the lead, insulation, and SAR boxes of a representative patient (ID12) at 3 T. A1: Closeup view of leads withing the high-resolution mesh regions. A2-A3: Schematic of mesh in the cubic area surrounding the lead tips and on the lead electrodes. A3: View of interconnections between electrodes and the central core. The coil’s input power was adjusted to generate a GHSAR of 3 W/kg. The distribution of 1g-SAR on a plane passing through the lead tips, the distribution of B_1_^+^ on an axial plane passing through the patient’s forehead, and the lead’s contacts’ geometrical details are also presented.

Figure 3 presents examples of the mesh density on the surface of leads, their insulation, and SAR boxes at 3 T for a representative patient with bilateral leads (ID12). Figure 3 also depicts the 1 g-SAR distribution on a plane passing through the lead tips and the B_1_^+^ distribution on an axial plane passing through the center of the head when the coil inputs are adjusted to maintain the GHSAR at 3 W/kg.

## 3 RESULTS

Figure 4 displays an example of the distribution of the 1 g-SAR on an axial plane passing through distal electrode contacts for each of the two exposure limits (patient ID12). The box plots presenting the 1 g-SAR_max_, mean of B_1_^+^, and mean of ‖**E**‖ for all lead models are provided in Figure 5 (B_1_^+^ and ‖**E**‖ are averaged over an axial plane passing through the center of the head shown in Figure 3). When the input power of the coils was adjusted to generate a GHSAR of 3 W/kg, the 1 g-SAR_max_ (mean ± standard deviation) was 104 ± 99 W/kg at 1.5 T, 32 ± 30 W/kg at 3 T, 19 ± 15 W/kg at 7 T, and 29 ± 17 W/kg at 10.5 T (Table S1). A one-way repeated measure analysis of variance (ANOVA) was conducted on lead models to examine the effects of different field strengths on RF power deposition near the lead tip. The results revealed that when the GHSAR was fixed, the variation of the field strength and resonant frequency led to statistically significant differences in power deposition near the lead tip (*F*(3, 96) = 24.09, *p* < 1.04 E-11). A post-hoc pairwise comparison test revealed that, for a fixed GHSAR, local SAR at 1.5 T was significantly higher than SAR at all other field strengths (*p* < .001 compared to 3, 7, and 10.5 T). Moreover, SAR at 3 T was significantly higher than that at 7 T (*p* = .008) but not significantly different from SAR at 10.5 T (*p* = .54). Finally, local SAR at 7 T was significantly lower than that at all other field strengths (*p* = .001 compared to10.5 T).

**Figure 4:**
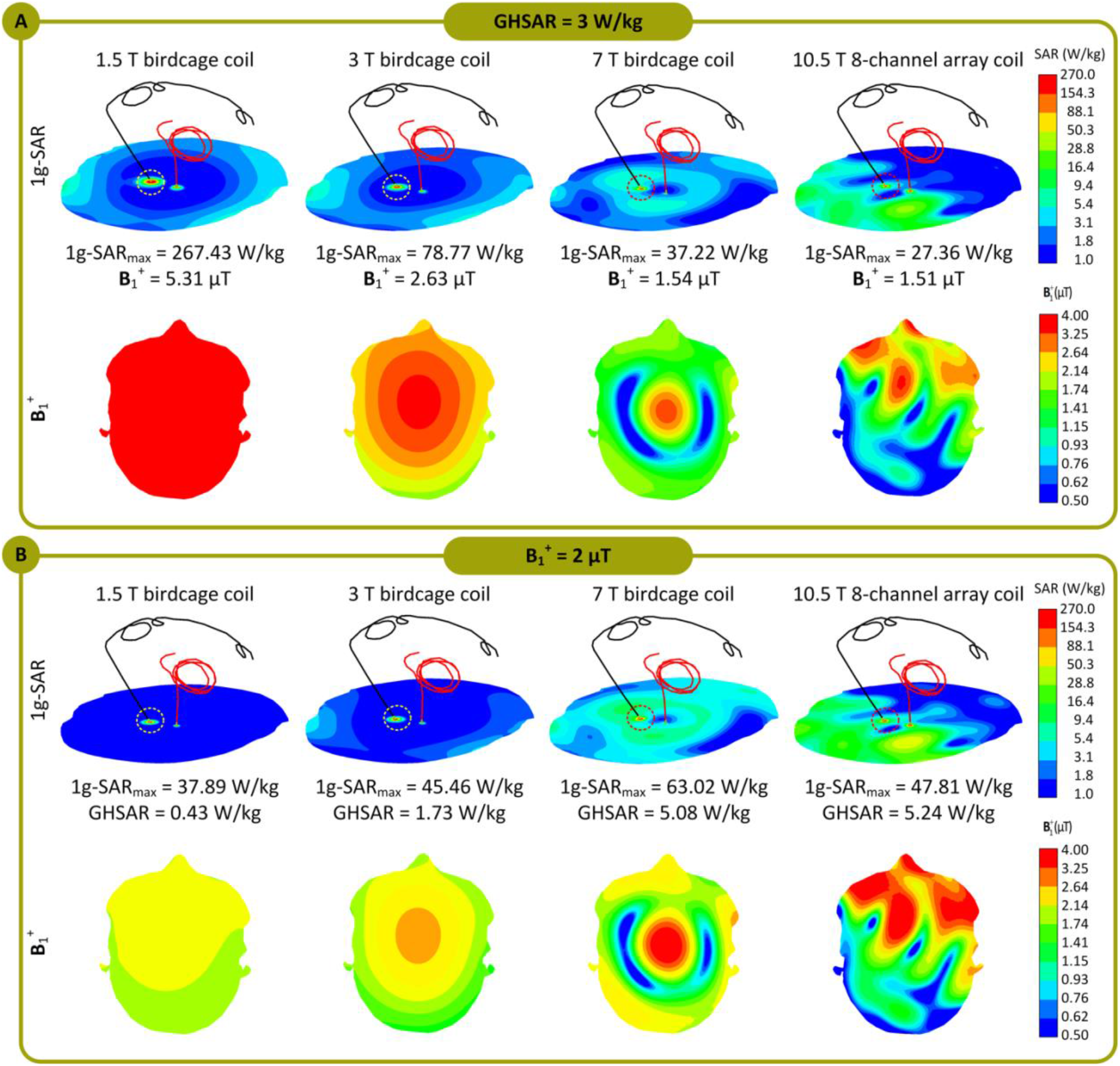
1g-SAR distributions in patient 12 (ID12) for the 1.5 T, 3 T, 7 T, and 10.5 T coils over an axial plane passing through the electrode contacts. The coils’ input powers are adjusted (A) to generate a GHSAR of 3 W/kg and (B) to generate an average of B_1_^**+**^ = 2 μT over an axial plane passing through the patient’s forehead. The B_1_^**+**^ field distributions over the mentioned axial plane, which passes through the patient’s forehead, are also presented. Additionally, 1g-SAR_max_, the average of B_1_^**+**^ field, and GHSAR are also reported for all four coils.

**Figure 5:**
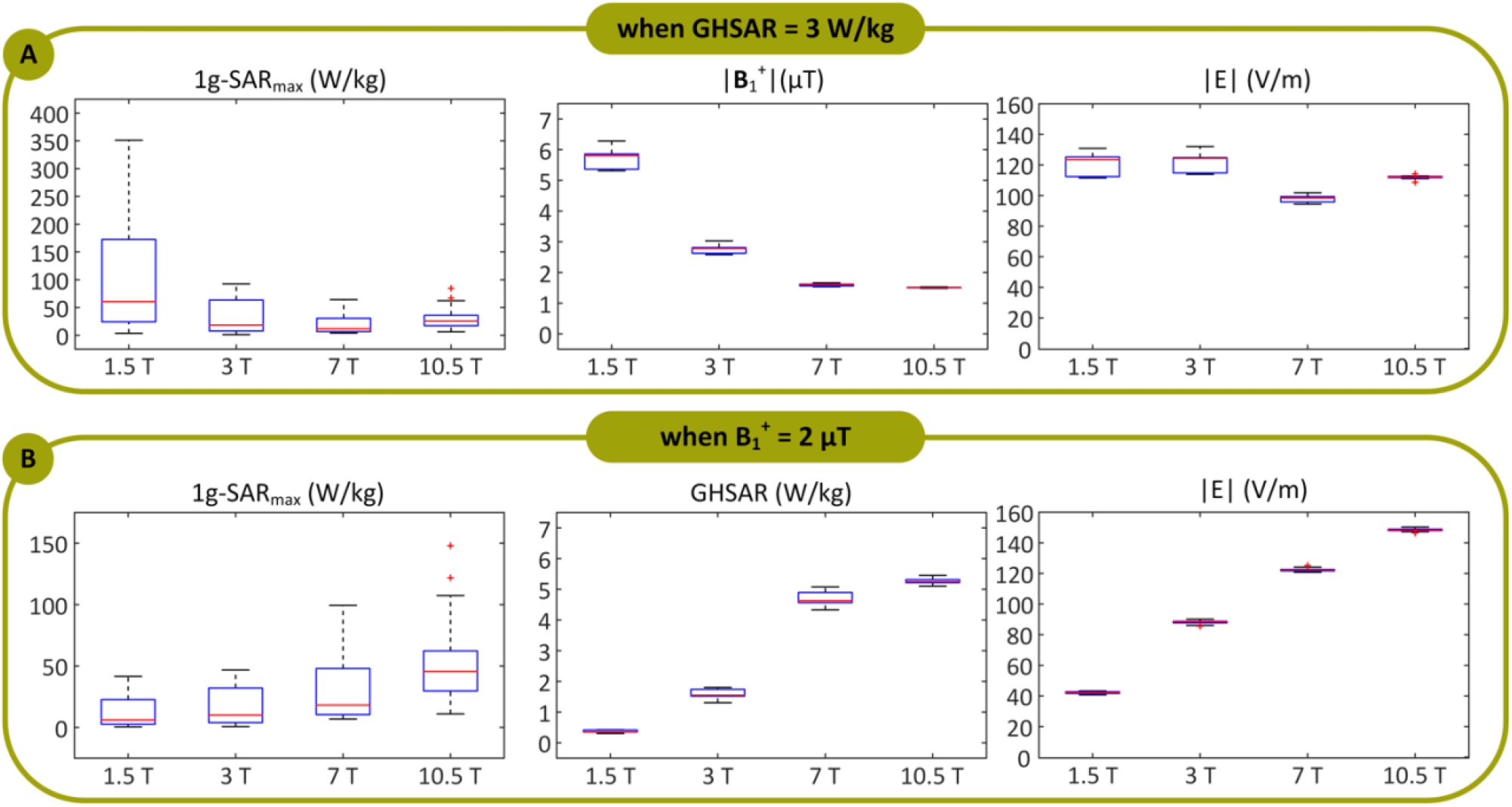
1g-SAR_max_, B_1_^+^, and ‖***E***‖ when the coils’ input powers were balanced to produce a global head SAR (GHSAR) of 3 W/kg, and (B) 1g-SAR_max_ and GHSAR when the coils’ input powers were adjusted to generate an average of B_1_^+^ = 2 μT, over all 33 DBS lead models for the 1.5 T, 3 T, 7 T, and 10.5 T coils. Box and whisker plots display the range, median, and interquartile range (IQR) of the data. Each outlier was specified with a red ‘+’ symbol.

When the output power was adjusted to generate a mean B_1_^+^ = 2 *µ*T at the center of the head, the 1 g-SAR_max_ was 13 ± 12 W/kg for the 1.5 T coil, 17 ± 16 W/kg for the 3 T coil, 29 ± 24 W/kg for the 7 T coil, and 51 ± 31 W/kg for the 10.5 T coil (Table S1). A one-way repeated measure ANOVA indicated that when the mean B_1_^+^ was fixed, the variation of the field strength/resonant frequency led to statistically significant differences in local SAR. A post-hoc pairwise comparison test confirmed a monotonous increase in the mean of local SAR as the field strength increased. For a fixed B_1_^+^, SAR was significantly lower at 1.5 T than that at all other field strengths (*p* = .02 compared to 3 T and *p* < .001 compared to 7 T and 10.5 T). In addition, SAR at 3 T was significantly lower than that at 7 T (*p* = .003) and 10.5 T (*p* < .001), and SAR at 7 T was significantly lower than that at 10.5 T (*p* < .001).

## 4 DISCUSSION AND CONCLUSIONS

The market for active implantable medical devices is increasing rapidly due to the aging population and rising prevalence of chronic diseases.[40] As MRI is now the standard imaging modality for various neurological, cardiovascular, and musculoskeletal disorders, the instances in which patients with electronic implants are indicated for MRI are also rapidly increasing. Currently, RF heating caused by the interaction between MRI RF fields and the conductive leads of active implants is the major safety concern preventing patients with electronic implants from accessing MRI. The issue is a complicated multivariable problem involving several interplaying factors, including the implant configuration and position within the patient’s body [16, 41, 42] and anatomy and the position inside the RF coil [38, 43, 44], the orientation and phase of the incident electric fields [35, 45-47], and the RF transmit operating frequency [6, 39].

Over the past two decades, numerous studies have investigated RF heating of elongated implants during MRI at mid-field strength, namely 1.5 T. In contrast, scarce data exist on how increasing the magnetic field strength and, by proxy, the resonant frequency affects implant heating. We conducted a systematic simulation study to compare local SAR near DBS lead tips in a range of resonant frequencies corresponding to 1.5 T to 10.5 T MRI. Historically, MR-Conditional labels of implantable devices have imposed limitations on the global SAR, which could be either the head SAR in the case of brain implants or the total body SAR in the case of implants in the torso and abdomen. Selection of global SAR as outcome measure was primarily because the global SAR was (and still is) automatically calculated and reported by all scanners. The first MR-Conditional guidelines for DBS devices, for example, limited the GHSAR to 0.1 W/kg instead of the 3.2 W/kg, which is the FDA recommended limit for scanning in the absence of implants [48]. Recent guidelines offer the option to cap either B_1_^+^ (for newer scanners that report B_1_^+^rms) or the global SAR as previously indicated. However, although calculation of B_1_^+^ may be straightforward at lower frequencies where the field is spatially homogeneous, there are open questions as how to quantify RMS B_1_^+^ at field strengths ≥ 7 T where the RF field is substantially less homogenous. The issue is even more pressing in the presence of implants, as they can affect the coil’s loading much more severely at higher fields. Figure 6 gives the B_1_^+^ maps of all four coils in a head model without any implant (top row), and in a head model with a DBS lead (bottom row). As it can be observed, the field becomes increasingly more inhomogeneous at higher fields when the implant is present.

**Figure 6:**
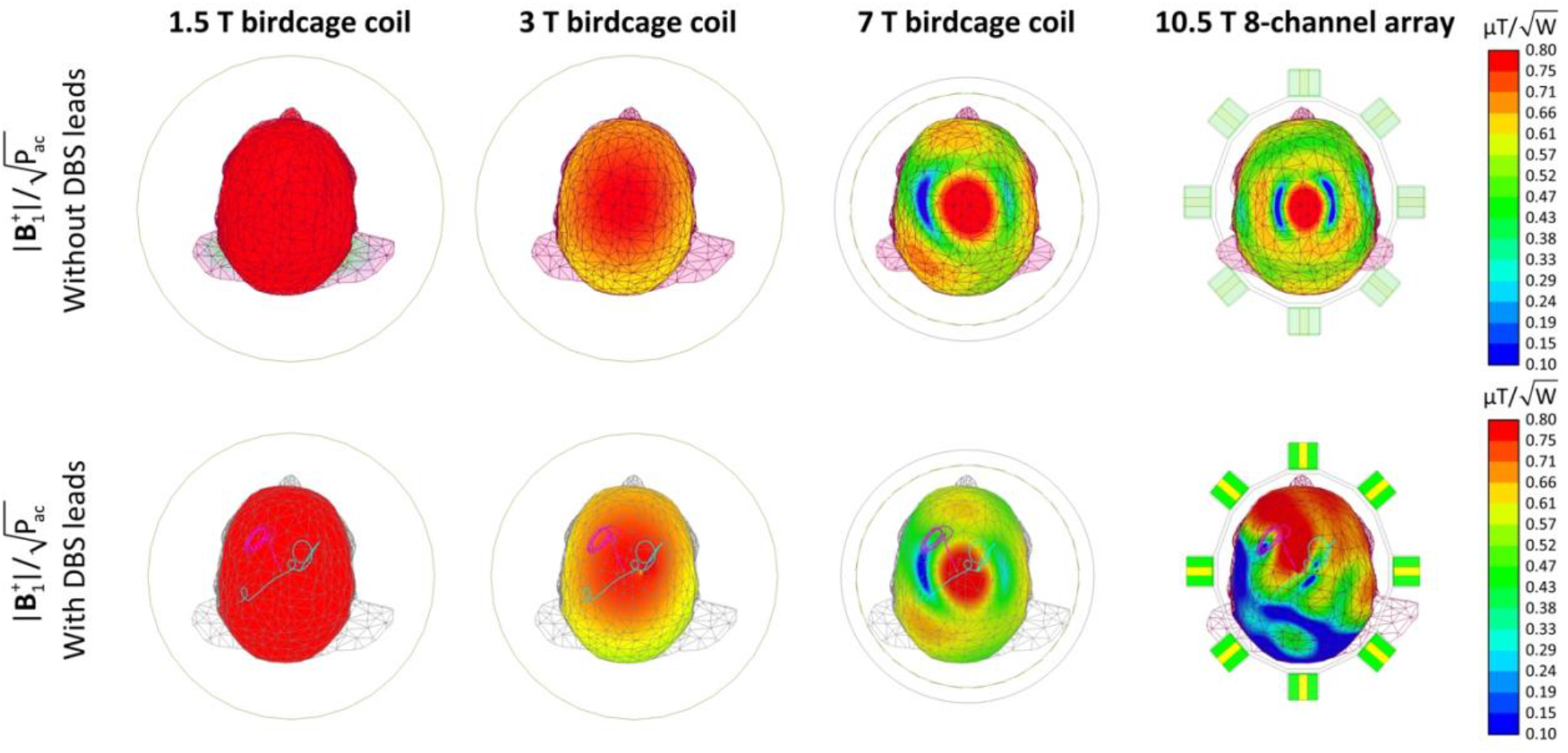
Normalized B_1_^+^ maps of the 1.5 T, 3 T, 7 T and 10.5 T coils on an axial plane passing through the iso-center in human head model with no implants (top rows) and with implants (bottom row). All four coils are driven in the CP excitation mode, where their inputs are adjusted to keep the accepted power (Pac) as 1 W. The B_1_^+^ maps of the second row were acquired for patient 12 (ID12).

In general, imposing the global SAR limit is more conservative. Specifically, Figure 5 demonstrates that one needs to settle for a lower B_1_^+^ magnitude at higher resonant frequencies to adhere to the GHSAR limit. This result is predictable, considering that the SAR is proportional to ‖*E*‖^2^ (see Equation 1) and that Maxwell’s equations state the following:

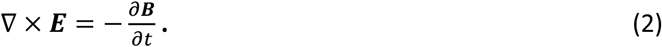

Therefore, capping the GHSAR keeps the magnitude of the **E** field relatively constant over the range of resonant frequencies by reducing the magnitude of B_1_^+^ at higher frequencies. The local RF heating generated at the lead tips is due to the current induced in the lead wires due to capacitive coupling with the **E** field and the inductive coupling with B_1_^+^. Therefore, for a constant GHSAR, scanners at lower resonant frequencies tend to generate higher local heating at the tips of implanted leads. In contrast, when the magnitude of B_1_^+^ is constant over the range of resonant frequencies, coils with higher resonant frequencies generate a higher **E** field according to Equation 2, leading to a higher local SAR around the implanted leads. It should be noted, however, that when the length of the lead is close to the resonance length, which is *λ*/2 where *λ* is the wavelength of the RF wave in the tissue, excessive heating could occur due to the resonance phenomenon [49, 50]. In this work we only examined 40 cm leads and thus the results should not be interpolated to other lead lengths.

This work has several limitations. First, we have used simplified DBS lead geometries with a straight internal core, as opposed to clinically available leads which have multiple helical wires each connecting to one electrode contact. Leads with helical cores will have different electric length than their apparent length, that is, a helical lead with a 40-cm apparent length may have substantially longer internal wires. It is hard to isolate the effect of the length as it also affects the trajectory (i.e., leads with different apparent length inevitably follow different trajectories), as well as the phenomenology of coupling with electric field (i.e., leads with helical interwire of different pitch with have different distributed inductance that could affect their resonance length). More study is needed with leads of different length and structure to better assess the effect of length. Furthermore, our models represent lead-only DBS systems. In a fully implanted device leads are connected to an extension cable (∼60 cm) which is then routed along the neck toward a pulse generator implanted in the chest. Although a fully implanted system will have a longer length, our simulation result of a heterogenous dispersive body model implanted with a full DBS system showed a similar trend in SAR and B_1_^+^ (See Supporting Information S2). Also, we did not correct for B_1_^+^ field inhomogeneity at field strengths above 7 T which may happen in practice. Finally, low, and ultra-low scanners (e.g., at 0.55 T and 65 mT) are absent from the current study but should be considered in future investigations especially as they are more likely to be used for implant imaging. Regardless of these limitations, this work presents the first systematic study of effect of field strength on RF heating of elongated leads. Our results agree with recent studies that compared RF heating of DBS devices across 1.5 T and 3 T scanners [51, 52] and provide useful information for a comparative evaluation of RF risks across a wide range of field strengths to help selecting appropriate safety criteria.

## SUPPORTING INFORMATION

Additional Supporting Information can be found in the Supporting Information section.

## Notes

### Competing Interest Statement

The authors have declared no competing interest.

