## Supporting Information S1 for "Effect of field strength on RF power deposition near conductive leads: A simulation study of SAR in DBS lead models during MRI at 1.5 T - 10.5 T"

**Supporting Information Figure S1:**

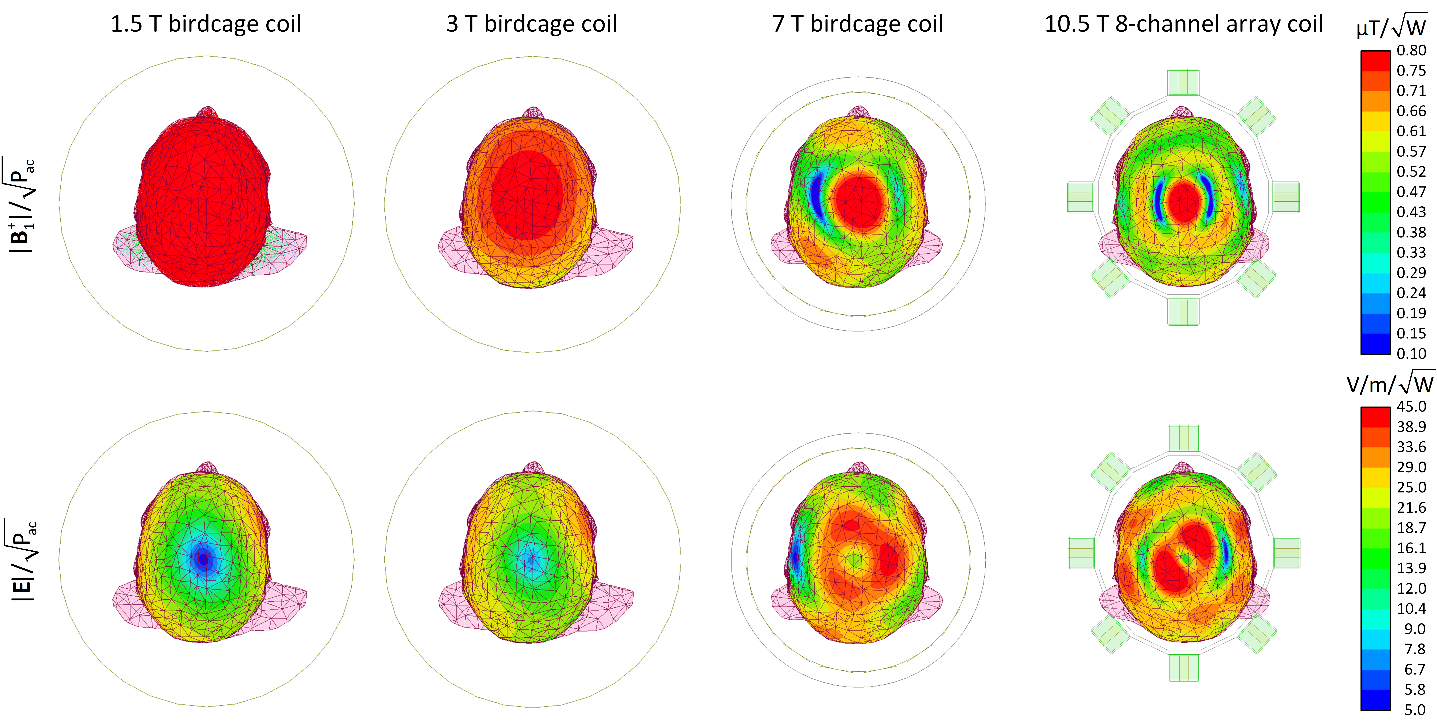

**Figure S1** - Normalized **B**_1_^+^ and **E** fields distributions of the 1.5 T, 3 T, 7 T and 10.5 T coils demonstrated in Figure 1 over the isocenter axial plane within a uniform human head model with no implants. All four coils are driven in the CP excitation mode, where their inputs are adjusted to keep the accepted power as 1 W.

**Supporting Information Table S1:**

**Table S1** – The maximum of 1g-averaged SAR

| Coil type | Field strength |  | when GHSAR = 3 W/kg | | when **B**_1_^+^ = 2 µT | |
| --- | --- | --- | --- | --- | --- | --- |
|  |  |  | 1g-SAR_max_ (W/kg) | **B**_1_^+^ (µT) | 1g-SAR_max_ (W/kg) | GHSAR (W/kg) |
| Birdcage coil | 1.5 T | Mean | 103.78 | 5.74 | 12.59 | 0.37 |
|  |  | SD | 98.51 | 0.33 | 12.14 | 0.04 |
| Birdcage coil | 3 T | Mean | 32.20 | 2.77 | 16.67 | 1.58 |
|  |  | SD | 29.67 | 0.15 | 15.35 | 0.16 |
| Birdcage coil | 7 T | Mean | 18.64 | 1.60 | 29.05 | 4.70 |
|  |  | SD | 15.23 | 0.04 | 23.86 | 0.22 |
| 8-channel array coil | 10.5 T | Mean | 29.13 | 1.51 | 51.04 | 5.27 |
|  |  | SD | 17.69 | 0.01 | 31.11 | 0.09 |
