## Supporting Information S2 for "Effect of field strength on RF power deposition near conductive leads: A simulation study of SAR in DBS lead models during MRI at 1.5 T - 10.5 T"

**Supplementary Information S2**

We simulated a detailed body model consisting of 21 tissues that expanded to the chest, with dielectric properties varying by frequency. Tissue properties at each frequency were retrieved from IT’IS Foundation website (https://itis.swiss/virtual-population/tissue-properties/database/dielectric-properties/) (see table below for values).

| Tissue clusters | Mass  density (Kg/m^3^) | **1.5 T (64 MHz)** | | **3 T (127 MHz)** | | **7 T (297 MHz)** | | **10.5 T (447 MHz)** | |
| --- | --- | --- | --- | --- | --- | --- | --- | --- | --- |
|  |  | σ (S/m) | ε | σ (S/m) | ε | σ (S/m) | ε | σ (S/m) | ε |
| Air | 1 | 0.00E+0 | 1.00E+0 | 0.00E+0 | 1.00E+0 | 0.00E+0 | 1.00E+0 | 0.00E+0 | 1.00E+0 |
| Bone Marrow (Yellow) | 980 | 2.16E-2 | 7.21E+0 | 2.41E-2 | 6.24E+0 | 2.78E-2 | 5.76E+0 | 3.07E-2 | 5.64E+0 |
| Cerebrospinal Fluid | 1007 | 2.07E+0 | 9.73E+1 | 2.14E+0 | 8.42E+1 | 2.22E+0 | 7.28E+1 | 2.26E+0 | 7.05E+1 |
| Blood | 1050 | 1.21E+0 | 8.64E+1 | 1.25E+0 | 7.33E+1 | 1.32E+0 | 6.57E+1 | 1.37E+0 | 6.37E+1 |
| Bone (Cancellous) | 1178 | 1.61E-1 | 3.09E+1 | 1.80E-1 | 2.63E+1 | 2.15E-1 | 2.32E+1 | 2.44E-1 | 2.22E+1 |
| Bone (Cortical) | 1908 | 5.95E-2 | 1.67E+1 | 6.72E-2 | 1.47E+1 | 8.24E-2 | 1.35E+1 | 9.56E-2 | 1.30E+1 |
| Cartilage | 1100 | 4.52E-1 | 6.29E+1 | 4.88E-1 | 5.30E+1 | 5.51E-1 | 4.68E+1 | 6.03E-1 | 4.50E+1 |
| Cerebellum | 1045 | 7.19E-1 | 1.16E+2 | 8.28E-1 | 8.00E+1 | 9.71E-1 | 5.99E+1 | 1.05E+0 | 5.48E+1 |
| Muscle | 1090 | 6.88E-1 | 7.22E+1 | 7.19E-1 | 6.36E+1 | 7.70E-1 | 5.82E+1 | 8.08E-1 | 5.68E+1 |
| Spinal Cord | 1075 | 3.12E-1 | 5.51E+1 | 3.53E-1 | 4.42E+1 | 4.17E-1 | 3.70E+1 | 4.59E-1 | 3.49E+1 |
| Trachea | 1080 | 5.28E-1 | 5.89E+1 | 5.59E-1 | 5.06E+1 | 6.10E-1 | 4.53E+1 | 6.48E-1 | 4.38E+1 |
| Brain (Grey Matter) | 1045 | 5.11E-1 | 9.74E+1 | 5.86E-1 | 7.37E+1 | 6.91E-1 | 6.01E+1 | 7.57E-1 | 5.66E+1 |
| Brain (White Matter) | 1041 | 2.92E-1 | 6.78E+1 | 3.42E-1 | 5.27E+1 | 4.12E-1 | 4.38E+1 | 4.59E-1 | 4.15E+1 |
| Eye (Vitreous Humor) | 1005 | 1.50E+0 | 6.91E+1 | 1.51E+0 | 6.91E+1 | 1.52E+0 | 6.90E+1 | 1.54E+0 | 6.90E+1 |
| Lung (Inflated) | 394 | 2.89E-1 | 3.71E+1 | 3.15E-1 | 2.95E+1 | 3.56E-1 | 2.48E+1 | 3.82E-1 | 2.35E+1 |
| Nerve | 1075 | 3.12E-1 | 5.51E+1 | 3.53E-1 | 4.42E+1 | 4.17E-1 | 3.70E+1 | 4.59E-1 | 3.49E+1 |
| Esophagus | 1040 | 8.78E-1 | 8.58E+1 | 9.12E-1 | 7.50E+1 | 9.71E-1 | 6.88E+1 | 1.02E+0 | 6.71E+1 |
| Fat | 911 | 6.62E-2 | 1.36E+1 | 6.97E-2 | 1.24E+1 | 7.64E-2 | 1.17E+1 | 8.28E-2 | 1.16E+1 |
| Eye (Retina) | 1036 | 5.11E-1 | 9.74E+1 | 5.86E-1 | 7.37E+1 | 6.91E-1 | 6.01E+1 | 7.57E-1 | 5.66E+1 |
| SAT (Subcutaneous Fat) | 911 | 6.62E-2 | 1.36E+1 | 6.97E-2 | 1.24E+1 | 7.64E-2 | 1.17E+1 | 8.28E-2 | 1.16E+1 |
| Tissue (Average) | 1090 | 3.60E-1 | 6.40E+1 | 4.12E-1 | 3.97E+1 | 4.87E-1 | 3.80E+1 | 5.54E-1 | 3.65E+1 |

We positioned two DBS leads (ID #4) in the body model and calculated SAR under different RF exposure conditions.

We observed the same trend in SAR, although absolute SAR values varied between simplified and detailed models. The simulation time, however, was substantially different (1820 minutes in average for the detailed model vs. 740 minutes for the simplified model). Therefore, we decided to run mass simulations with simplified head model.

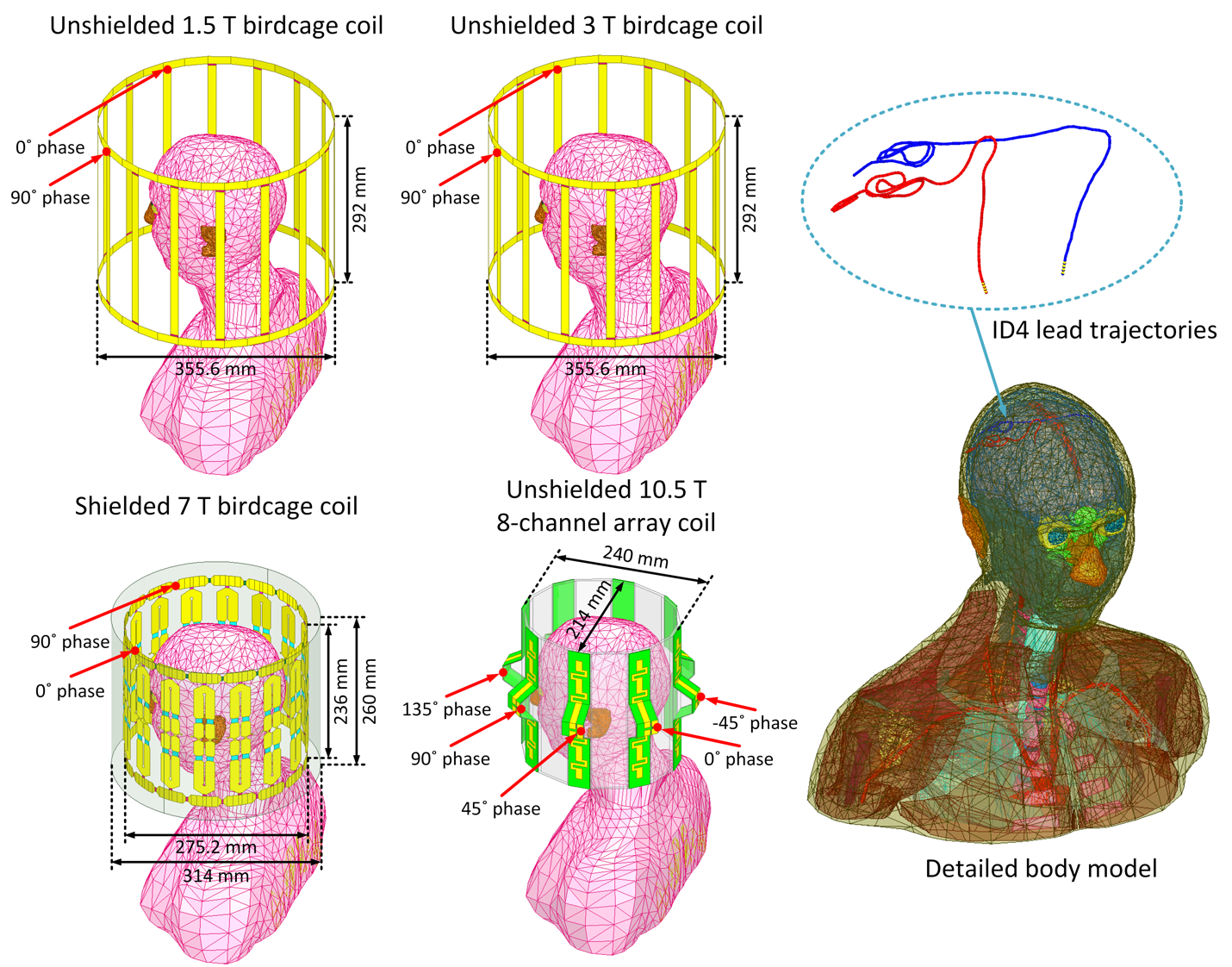

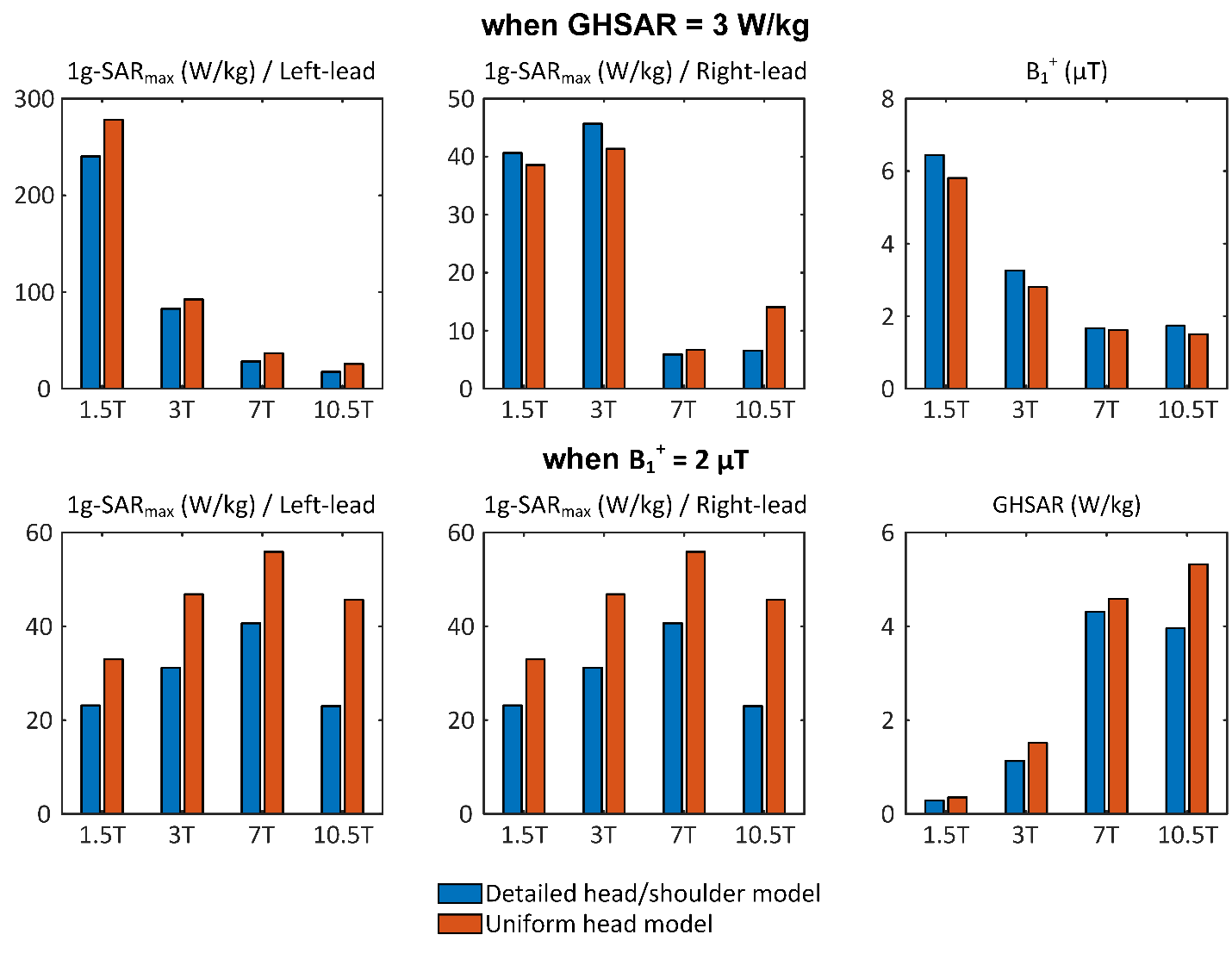
